# Not only age affects cardiovascular parameters, salivary biomarkers, and their correlation, but the level of physical conditioning changes this behavior in the elderly

**DOI:** 10.1101/2023.02.24.529896

**Authors:** Phabloo José de Venâncio de Camargos, Allisson Benatti Justino, Luis Carlos Oliveira Gonçalves, Rafael Joviano Souza de Barros, Miguel Junior Sordi Bortolini, Foued Salmen Espindola, Aníbal Monteiro de Magalhães Neto

## Abstract

To compare the physical conditioning, hemodynamic, and salivary biomarkers between elderly athletes and the physically active elderly. 14 men: EA (n = 8) and PAE (n = 6). Collection times (T0; TE; T5; T15). A negative correlation was found between SF and cardiovascular parameters, BL, and STP in both groups, but this was almost double among PAE. For HR and SBP, there was a faster recovery in EA. The EA increase was correlated with SBP, while for PAE it was correlated with HR. BL showed an increase in TE, reaching 481% in EA and 639% in PAE. SNO showed a similar increase for the groups at the TE, but at T5, while EA already showed a reduction, PAE saw a 94% increase, with a slower decay for this group at T15. The SF presented the negative Δ% was almost double in PAE, with a quick recovery already at T5 for EA and levels still negative at all times for PAE. For SIgA-s, there was an increase of 37% in EA and only 7% at the TE in PAE; 41% in EA and 15% in PAE for T5; and 26% in EA and 14% in PAE at T15. SA showed a higher peak in EA (TE) and less acute in PAE (T5) but there was a decrease among both at T15. STP increased by 126% in EA and 438% in PAE, already showing a return at T5 for EA, but increasing by 213% in PAE. Negative levels were reached at T15 for EA but levels remained high in PAE. Levels of physical conditioning affect cardiovascular parameters, salivary biomarkers, and their correlation within the over-60s.

## 1. Introduction

It is widely discussed by the scientific community that sarcopenia generated by physical inactivity added to mental depression over the years can affect the quality of life of the elderly.^1^

This makes clear the need for regular exercise, minimizing the loss of bone and muscle mass, providing better balance, and reducing the chances of falls, representing a significant problem at this age.^2^

Therefore, using a sport method that reproduces natural conditions of the sport practiced in a controlled and intentional way aims to understand the immunometabolic differences generated in different populations. In the case of older people who regularly exercise but with different levels of physical conditioning, it is essential. for the knowledge of geriatricians and gerontologists.^3-5^

The main objective of this work was to evaluate and compare the physical conditioning, hemodynamic, blood, and salivary biomarkers between elderly considered active and athletes before, at the end, and during the recovery period of an exercise test on a treadmill.

## 2. Method

### 2.1 Participants

Thirty men over the age of 60, each of whom were assiduous competitors and regular exercisers with a minimum experience of 20 years in street racing events, were selected. They were first examined by a general dentist who used the Plaque Control Index (PCI), Bleeding Index (SI), and Simplified Oral Hygiene Index (OHIS) and checked for xerostomia symptoms. The participants in good oral health moved on to the next phase of the experiment. Those who were not were treated by the same dentist but did not take any further part in the study.

The selected individuals underwent a careful anamnesis of general health conditions, training routine, and competitions. The final sample of elderly people comprised 14 men divided into two groups: elderly athletes (EA: n = 8) and the physically active elderly (PAE: n = 6).

The subjects’ characteristics were as follows: **EA** 63.1 ± 6.2 years, weight 68.4 ± 8.1 kg, height 170.1 ± 1.9 cm, body mass index 20.5 ± 1.0 kg/m2, resting heart rate 67.7 ± 7.0 bpm, maximal heart rate 154.7 ± 5.2 bpm, VO2 max 32.8 ± 5.5 ml/kg/min). **PAE** 63.8 ± 4.0 years, weight 70.4 ± 3.1 kg, height 168.5 ± 2.8 cm, body mass index 23.5 ± 1.5 kg/m2, resting heart rate 76.7 ± 1.5 bpm, maximal heart rate 147.0 ± 10.2 bpm, VO2 max 24.4 ± 7.0 ml/kg/min).

### 2.2 Experimental Design

In the 48 hours before the exercise test, the participants did not perform any physical exercise. They ate their last meal at least two hours before the test. They were instructed to thoroughly clean their mouths and drink water ad libitum the day before the test. No licit or illicit stimulant substances (i.e., coffee, guarana, teas, and other xanthines, thermogenic, or central nervous system stimulants) or anything containing dyes could be ingested during this period.

For adaptation purposes, all participants had to perform two training sessions on different days. The stress test was carried out on a treadmill (Ecafix®, Brazil). The original Bruce protocol was used. Heart rate was monitored throughout the exercise test by 12-lead electrocardiogram using Ergo PC 13 for Windows. Systolic blood pressure (SBP) and diastolic blood pressure (DBP) were measured using the auscultatory method, with a sphygmomanometer and stethoscope (Tycos®, USA): ([2 × diastolic pressure] + systolic pressure)/3.

### 2.3 Procedures and Measurements

Salivation was stimulated by chewing gum (Cadbury Adams Brasil Ind®) weighing 1.5g. Chewing took place naturally and personally without instructions on speed, strength, and frequency, though the participants were instructed to chew the gum on only one side of the mouth. On the other side, the Tubo Salivette® device was used for quick and safe saliva collection. The participants were given a chewing gum tablet to carry during street training sessions.

Saliva was placed in pre-cooled (4°C) mini-tubes. All samples were processed, and immediately after collection, centrifuged at 12,000 g; the pellet was discarded and the supernatant was frozen at 20ºC until the day of the analysis.

The stress test was performed on a treadmill (Ecafix®, Brazil). The original Bruce protocol was used. ^6^ Heart rate was monitored throughout the exercise test by 12-lead electrocardiogram using Ergo PC 13 for Windows. Systolic blood pressure (SBP) and diastolic blood pressure (DBP) were measured using the auscultatory method, with a sphygmomanometer and stethoscope (Tycos®, USA): ([2 × diastolic pressure] + systolic pressure)/3. All the above-mentioned physiological parameters were evaluated at rest (T0), during the exercise test, at the maximum peak of the exercise (TE), and on the stands five and 15 minutes after the end of the test (T5 and T15). All stress tests were performed in a private cardiology clinic in the presence of a physician. All participants had to perform two training sessions on different days to adapt. The exercise test was interrupted when any of these criteria were identified: elevation of diastolic blood pressure (DBP) > 120mm/Hg in normotensive individuals and > 140mm/Hg in primary hypertensive individuals; elevation of systolic blood pressure (SBP) > 260mm/Hg; a fall sustained SBP; clinical manifestations of typical severe chest pain; ST-segment depression > 3mm; ST-segment elevation > 2 mm in the lead without a q wave; complex ventricular arrhythmia; sustained supraventricular tachycardia onset; atrial tachycardia; atrial fibrillation; atrial block (second and third degree ventricular); signs of left ventricular failure; and the failure of monitoring and recording systems ^7^.

Total salivary protein was measured by the biuret method using a standard laboratory kit (UCFS DIASYS® cat. n° 1 0210 99 10 021 Germany) through two spectrophotometric readings, one primary at 604 nm and the other at 700 nm (Autoanalyser Architeet c8000, Abbot®, IL, USA).

The analysis of alpha-amylase activity was performed by a kinetic assay using the CNPG kit (Pro Biotec, Ind. Com. Diagnóstico para Saúde, Uberlândia, MG, Brazil). Assays were performed at room temperature (28°C) on microplates with ten microL of saliva diluted 100 times in saline and 300 microL of CNPG buffer. The microplates were read at 405 nm on a microplate reader (Amershan Biosciences, GE, Upsala, Sweden) programmed to give the results unit of salivary amylase activity/mL of saliva (U/mL). Total salivary protein was measured by the biuret method according to a standard laboratory kit (UCFS DIASYS® cat. n° 1 0210 99 10 021 Germany) through two spectrophotometric readings, one primary at 604nm and the other at 700nm (Autoanalyser Architeet c8000, Abbot ®, IL, USA).

The analysis of alpha-amylase activity was performed by a kinetic assay using the CNPG kit (Pro Biotec, Ind. Com. Diagnóstico para Saúde, Uberlândia, MG, Brazil). Assays were performed at room temperature (28°C) in microplates with ten microL of saliva diluted 100 times in saline and 300 microL of CNPG buffer. The microplates were read at 405nm on a microplate reader (Amershan Biosciences, GE, Upsala. Sweden) programmed to give the results unit of salivary amylase activity/mL of saliva (U/mL).

Total salivary protein was measured by the biuret method according to a standard laboratory kit (UCFS DIASYS® cat. n° 1 0210 99 10 021 Germany) through two spectrophotometric readings, one primary at 604nm and the other at 700nm (Autoanalyser Architeet c8000, Abbot ®, IL, USA).

The analysis of alpha-amylase activity was performed by a kinetic assay using the CNPG kit (Pro Biotec, Ind. Com. Diagnóstico para Saúde, Uberlândia, MG, Brazil). Assays were performed at room temperature (28°C) in microplates with ten microL of saliva diluted 100 times in saline and 300 microL of CNPG buffer. The microplates were read at 405nm on a microplate reader (Amershan Biosciences, GE, Upsala. Sweden) programmed to give the results unit of salivary amylase activity/mL of saliva (U/mL). of a 250 μM NaNO3 solution to verify the linearity of the reaction and calculate a conversion factor of absorbance values into concentration (line equation). The absorbance of the tests was quantified at 548 nm in a microplate spectrophotometer (Synergy Microplate Reader) with analysis by the BioTek Gen5 Data Analysis Software (BioTek instruments, Winooski, Vermont, USA).

The analysis of total IgA used the ELISA test according to laboratory routine. First, polystyrene plates (Maxi-Sorp, Nunc, Wohlen) were sensitized with anti-human IgA antibody (Sigma Chemical, Buchs), diluted at an optimal concentration in carbonate buffer, 0.06M (pH 9.6) for 12 hours at 4ºC. Then, the plates were washed and blocked with a specific buffer of sodium dihydrochloride.

Orthophenylenediamine (OPD). Saliva samples were diluted from 1:2 in 1% BSA-PBS-T and incubated for 1 hour at room temperature. After washing, biotinylated anti-IgA conjugate labeled with peroxidase diluted in the concentration to be used was added. The enzyme-substrate H2O2 + OPD (chromogen buffer) was incubated at room temperature for 1 hour. The results were expressed in ELISA indices (IE) for individual plate analysis. Optical density (OD) values were determined in an ELISA reader (Titertek Multiskan Plus, Flow Laboratories, USA) at 405 nm. After obtaining the results of total IgA, it was divided by the value of total protein in saliva, and the value of specific IgA was found.

Blood collection was performed in the left ear lobe of each volunteer. The first drop of blood was discarded to avoid contamination, with lactate eliminated in the sweat produced by the sweat glands. Then 25 microL of blood was collected in heparinized and calibrated capillaries. Blood lactate was analyzed by the electroenzymatic method.

### 2.4 Statistical Analysis

Initially, descriptive statistics were performed on the data, with measurements of position (mean, median, mode, and percentiles) and dispersion (amplitude, variance, standard deviation, and standard error).

Afterward, the univariate analysis of these data was performed using the Shapiro-Wilk normality test (because the sample was smaller than 30 individuals). The equal variance test would be applied if the Shapiro-Wilk test presented a result indicating normal distribution (P>0.05). For results with P>0.05, the paired T-Student test would follow; if P≤0.05, the paired T-Student test would follow the non-parametric Mann-Witney test. If the Shapiro-Wilk test presented a result indicating non-normal distribution (P≤0.05), the non-parametric Mann-Witney test would be applied directly.

Still, in the phase of the univariate analysis, the analysis of repeated measures ANOVA One Way dependent was performed because they were the same individuals in different conditions and moments.

So, for a better interpretation of the data, the individuals were divided into two groups according to their sex. Then, the calculation of percentage variation was applied:

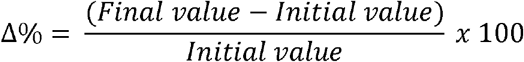

Cohen’s equations ^8^ were used to calculate the effect size for all variables to obtain Cohen d and r values

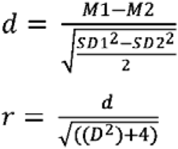

Where, M represent the means of observations and SD their respective standard deviations.Next, multivariate data analysis was performed using data mining and machine learning techniques.

In this phase, in order to seek a bivariate measure between the data, because the observations contain quantitative values, the Pearson and Spearman correlation tests were applied, with the Spearman correlation being used for a visual analysis using the heat map strategy and the Pearson test as an initial measure for the following machine learning analyses.

As exploratory models of machine learning: CLUSTER - Classical Clustering (Agglomerative Hierarchical Method) and Nearest neighbor (single linkage); ORDINATION – Principal component Analysis (PCA) and Correspondence Analysis (CA).

The Z score was previously applied because the observations contained non similar measurement units.

SigmaPlot 14.5 (Academic Perpetual License - Single User – ESD Systat® USA) and, Past 4.03 (Free version for Windows) were used to carry out the different statistical tests and produce the graphs. Finally, the correlation coefficients were presented using heat maps ^5^.

## 3. Results

A holistic and integrated analysis was carried out to avoid traditional dogmas and paradoxes.^5^ A Spearman rank correlation coefficient strategy was adopted so that the findings could be plotted on heat maps for better visualization.

A negative correlation was found between salivary flow and cardiovascular parameters, blood lactate, and total salivary proteins in both groups, but with correlation coefficients of almost double for the elderly only active the elderly athletes, as the heat maps show (Figs. 1 and 3).

**Figure 1.**
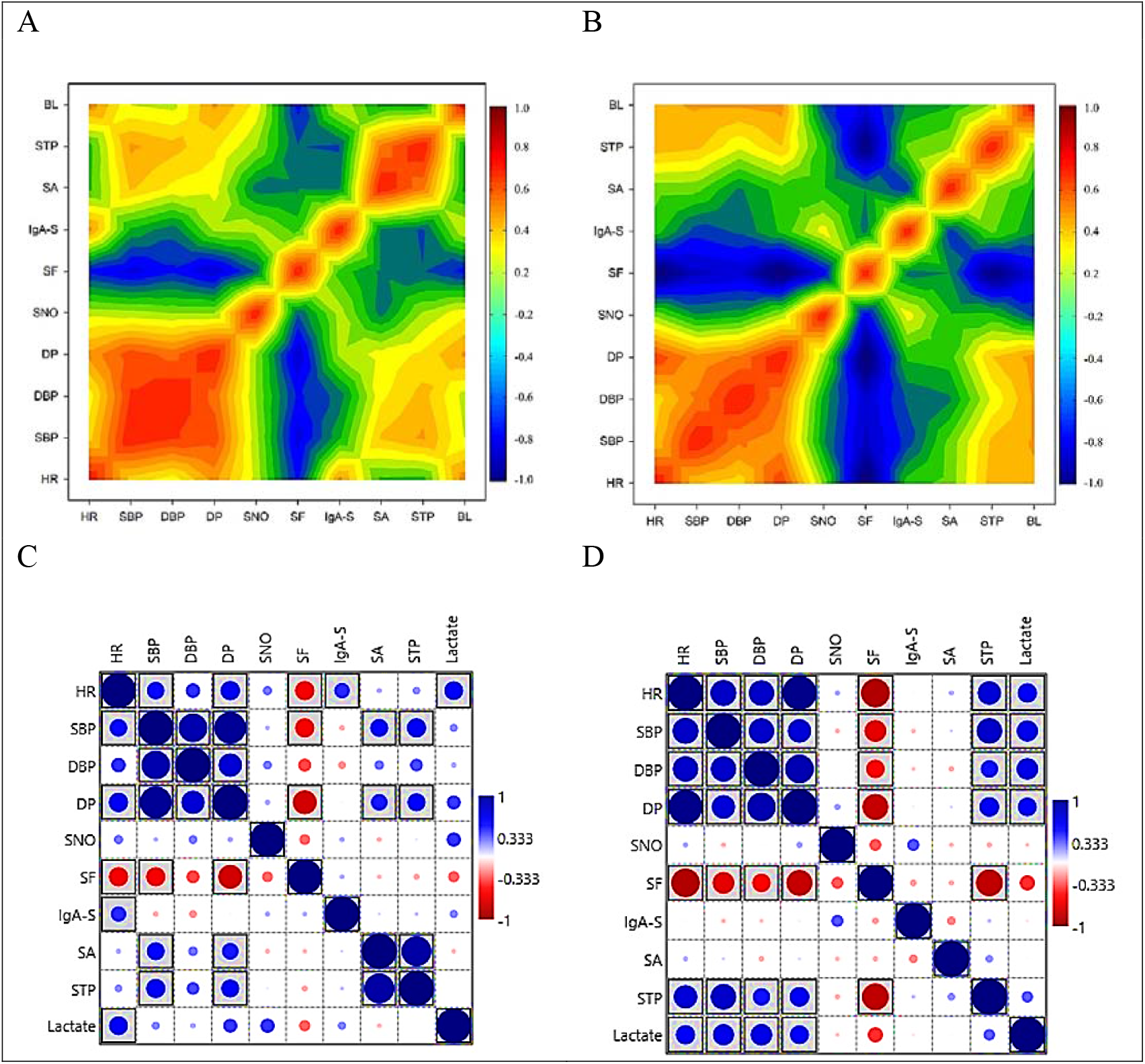
Heat map with holistic analysis (A and C – EA / B and D – PAE).

**Figure 2.**
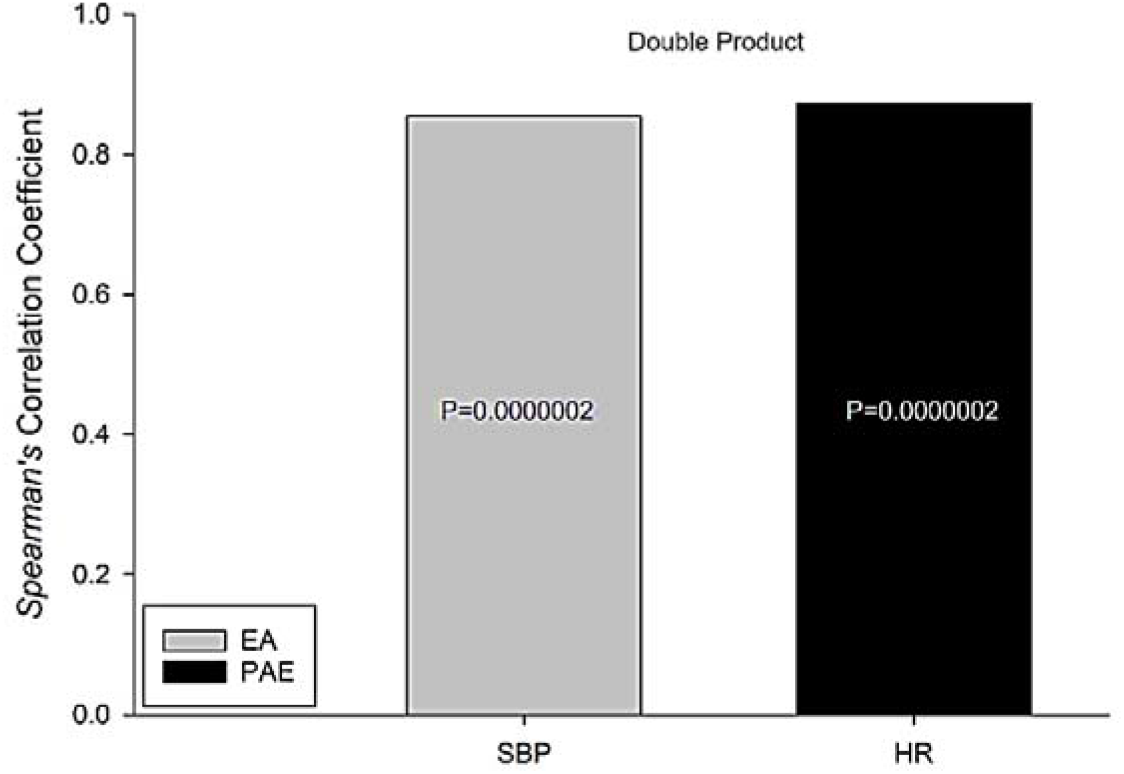
The double product of the elderly athletes is more closely correlated with systolic blood pressure, while the double product is more closely correlated with heart rate.

**Figure 3.**
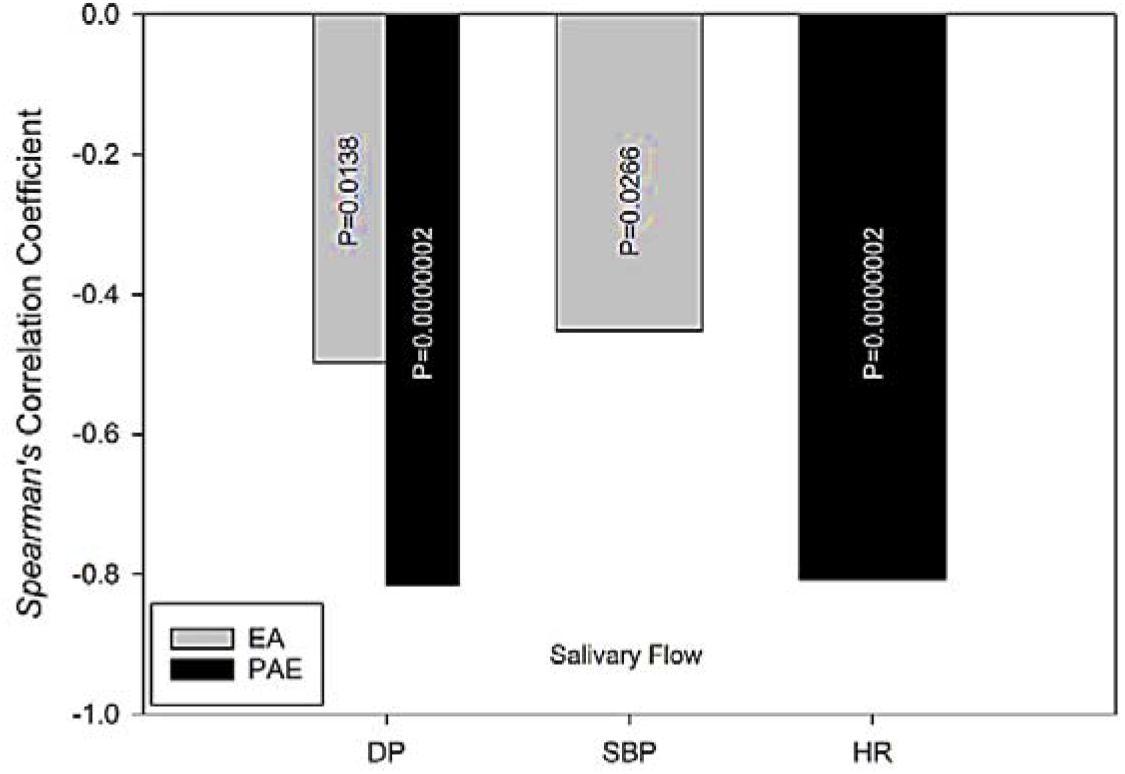
The salivary flow of both groups is negatively correlated with the double product. This correlation is almost double in the elderly athletes; in the first group it is caused by SBP and in the second by HR.

Also, to facilitate the visualization of the behavioral data in both groups, Table 2 presents the values of the size of the Cohen effect and the percentage variations concerning T0. As was expected, heart rate and systolic blood pressure recovered faster among EA. The opposite was the case for diastolic blood pressure and the double product.

**Table 1.** Values of effect size.

| Effect size | Small | Medium | Large |
| --- | --- | --- | --- |
| Cohen r | 0.10 | 0.30 | 0.50 |
| Cohen d | 0.20 | 0.50 | 0.80 |

Source: <sup>8</sup>

**Table 2.** The Cohen’s effect size (r) for significant salivary, blood, and cardiovascular biomarkers between groups and (the percentage variation) of T0.

| Parameters | Groups | TE | T5 | T15 |
| --- | --- | --- | --- | --- |
| Heart Rate | EA | 0.93 (125%) | 0.64 (27%) | 0.51 (-5%) |
|  | PAE | 0.99 (150%) | 0.79 (34%) | 0.79 (31%) |
| Systolic Blood Pressure | EA | 0.83 (53%) | 0.17 (5%) | 0.04 (1%) |
|  | PAE | 0.76 (54%) | 0.17 (-3%) | 0.68 (-15%) |
| Diastolic Blood Pressure | EA | 0.65 (26%) | 0.06 (2%) | 0.06 (-2%) |
|  | PAE | 0.69 (18%) | 0.13 (-2%) | 0.52 (-9%) |
| Double Product | EA | 0.91 (242%) | 0.57 (35%) | 0.40 (21%) |
|  | PAE | 0.96 (297%) | 0.60 (27%) | 0.22 (8%) |
| Salivary Nitric Oxide | EA | 0.16 (22%) | 0.08 (8%) | 0.03 (-3%) |
|  | PAE | 0.17 (34%) | 0.28 (94%) | 0.11 (27%) |
| Salivary Flow | EA | 0.62 (-28%) | 0.14 (5%) | 0.40 (14%) |
|  | PAE | 0.97 (-54%) | 0.93 (-38%) | 0.70 (-17%) |
| Salivary Immunoglobulin A | EA | 0.63 (37%) | 0.64 (41%) | 0.45 (26%) |
|  | PAE | 0.24 (7%) | 0.52 (15%) | 0.51 (14%) |
| Salivary Amylase | EA | 0.48 (130%) | 0.17 (26%) | 0.02 (3%) |
|  | PAE | 0.28 (68%) | 0.39 (100%) | 0.01 (2%) |
| Salivary Total Protein | EA | 0.49 (126%) | 0.02 (-1%) | 0.21 (-16%) |
|  | PAE | 0.78 (438%) | 0.59 (213%) | 0.37 (47%) |
| Blood Lactate | EA | 0.84 (481%) | 0.64 (398%) | 0.58 (353%) |
|  | PAE | 0.91 (639%) | 0.80 (495%) | 0.55 (196%) |

Regarding the latter double product, it was possible to observe an increase in both groups, but EA generated an intriguing finding: the increase was more closely correlated with the SBP (CC 0.854; *p* = 0.0000002), while for the PAE, the highest correlation was with HR (CC 0.872; *p* = 0.0000002; Figure 2).

As was expected, blood lactate showed an acute increase in time to exhaustion, reaching an increase of 481% in EA and 639% in PAE. However, what was not expected was the faster recovery of this metabolite in the PAE group, with a similar correlation between groups for DP and BL (Figure 4).

**Figure 4.**
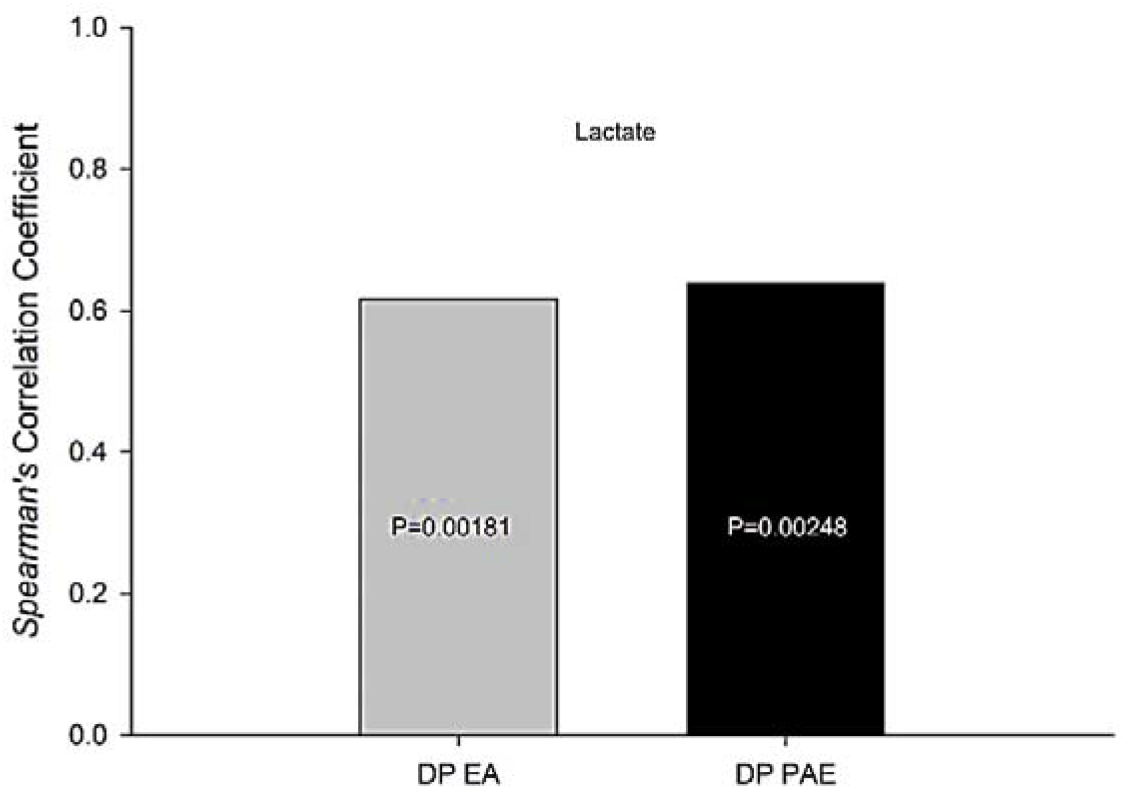
Blood lactate is positively correlated with double product in both groups.

The behavior of salivary biomarkers is shown in Figure 5 and Table 2 and their correlations with cardiovascular parameters in Figure 6, with very different designs.

**Figure 5.**
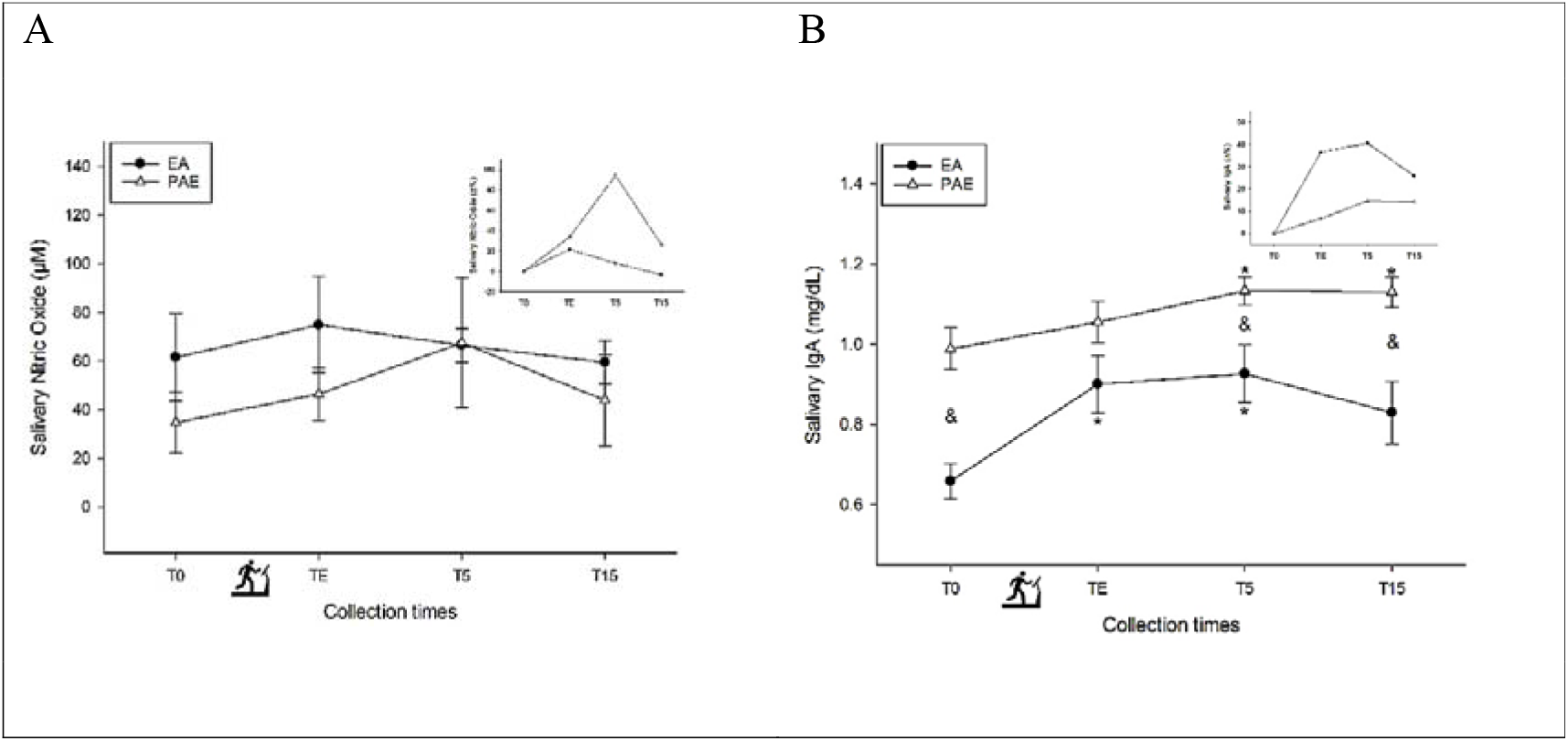
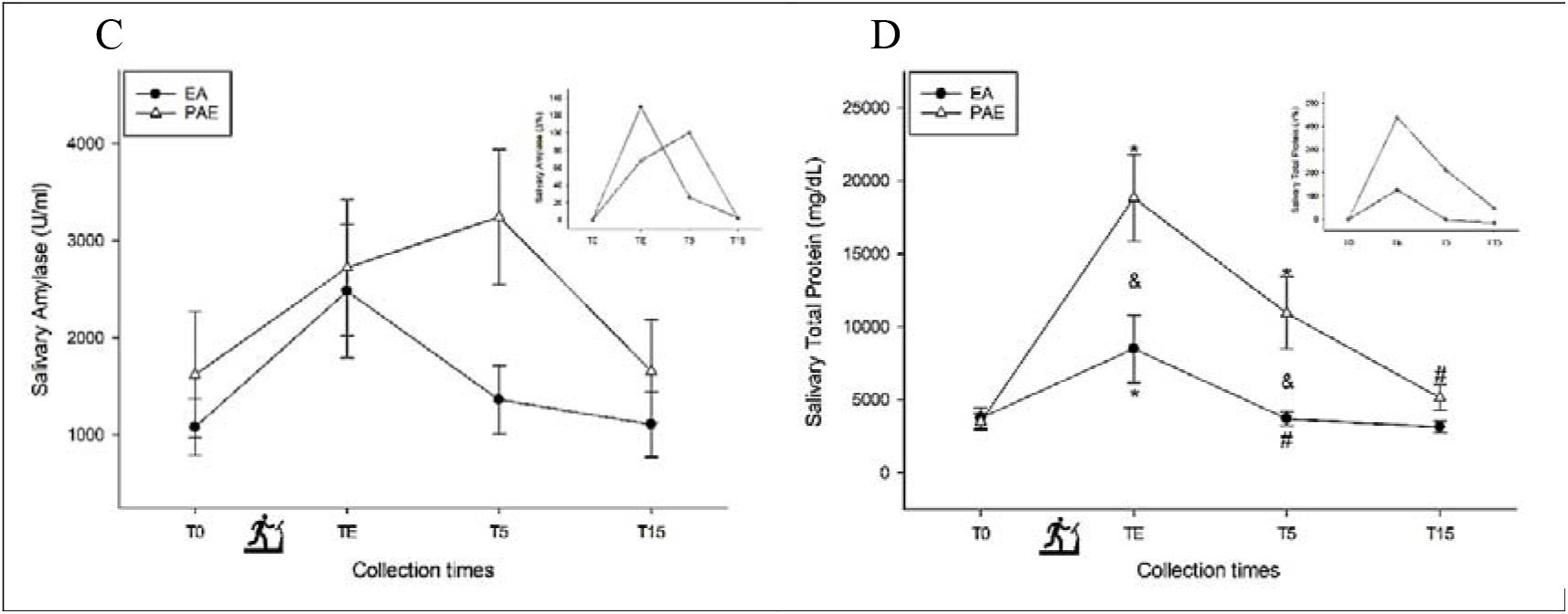
The behavior of salivary biomarkers (A – SNO; B – IgA-s; C – SA; D – STP).

**Figure 6.**
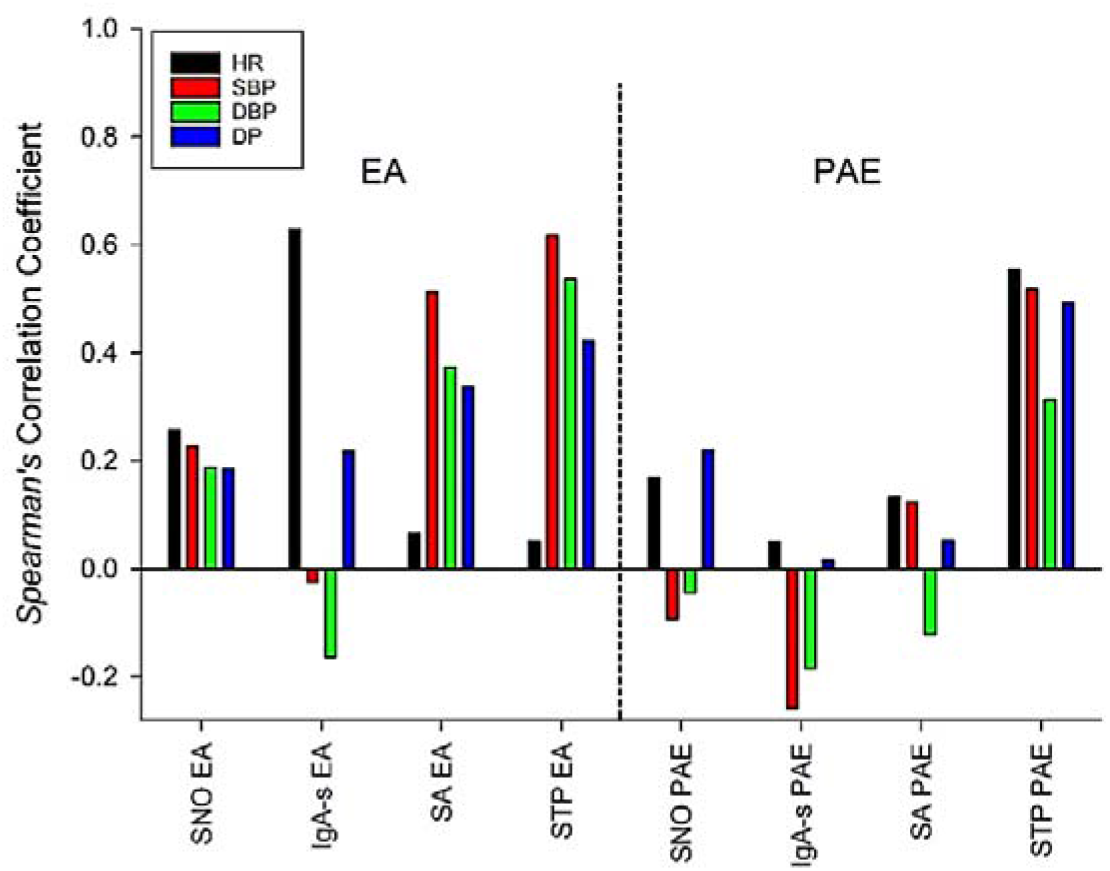
Correlation between salivary biomarkers and cardiovascular parameters.

Dendrogram with Euclidian distance of Salivary Biomarkers with nearest neighbor clustering plot: single linkage (Fig 7A) confirmed that the double product of the physically active group had a behavior mainly altered by heart rate, while in the group of athletes by systolic blood pressure. Regarding salivary proteins, the active group presented behavior more similar to diastolic blood pressure, while the group of athletes in the group of athletes was more similar to salivary amylase. The similarity between the lactate of both groups made it clear that the exercise intensity was high and similar, and network plot with Fruchterman-Reingold algorithm (Fig 7B) revealed that the salivary flow of only active individuals was highly dissimilar to the other biomarkers and that this flow negatively correlated with systolic blood pressure in the athletes. In contrast, in the physically active group, the negative correlation was greater with heart rate.

**Figure 7.**
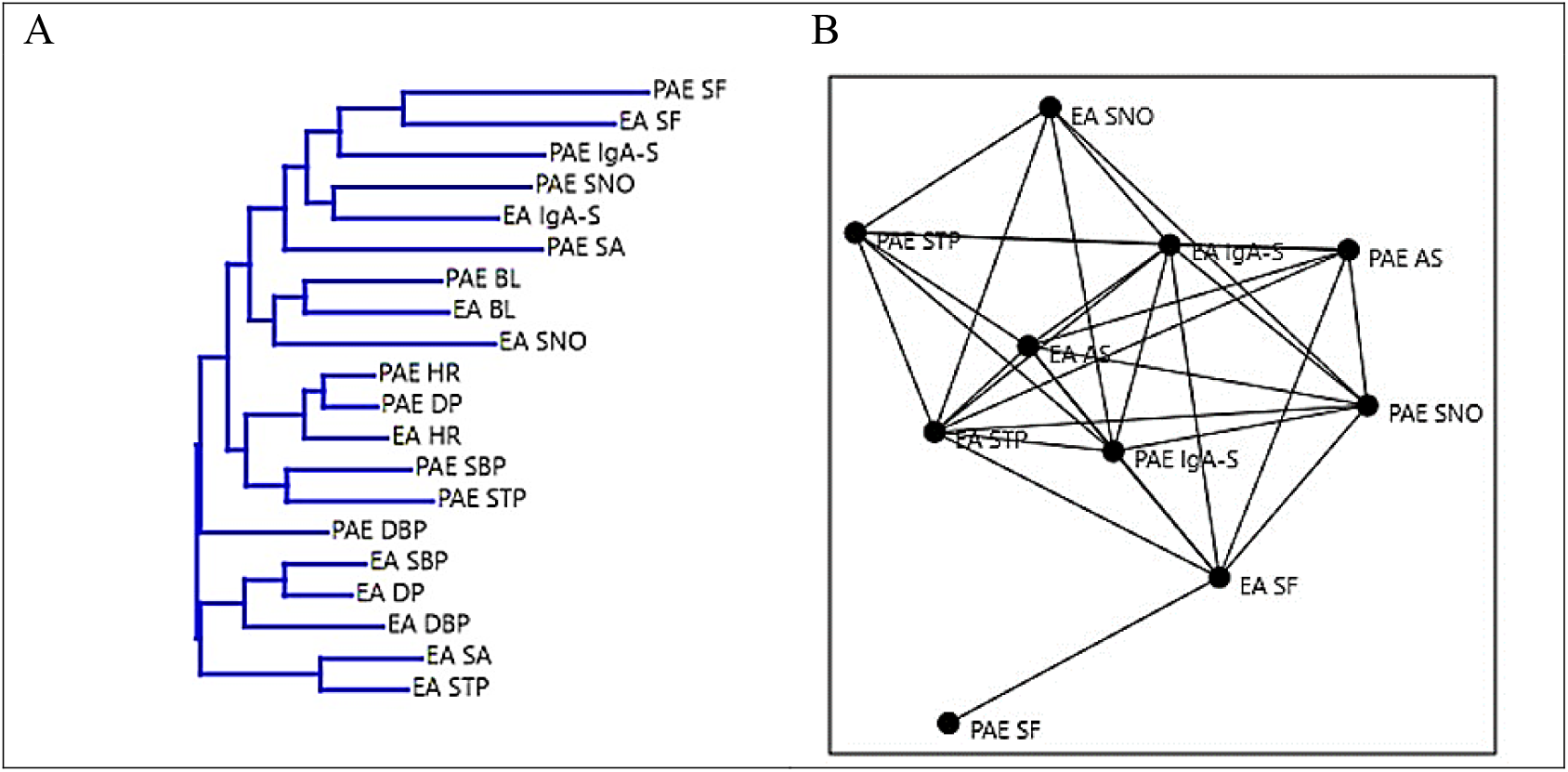
Dendrogram with Euclidian distance of Salivary Biomarkers. Nearest neighbor clustering plot: single linkage (Fig 7A), network plot with Fruchterman-Reingold algorithm (Fig 7B).

Salivary nitric oxide showed a similar increase in both groups at the time of exhaustion, but five minutes later (T5); while EA showed a reduction, PAE had a 94% increase, with a slower decay for the latter at T15.

Salivary flow was another biomarker where behaviour differed between the groups. The negative percentage variation was almost double in PAE. There was a rapid recovery at T5 for EA, and levels were negative at all times for PAE.

Salivary immunoglobulin A showed an increase of 37% in EA and only 7% in PAE (TE). They reached 41% in EA and 15% in PAE for T5 and 26% in EA and 14% in PAE for T15.

Salivary amylase showed a higher peak in the EA group (TE). It was less acute in PAE (T5) but decreased to baseline values for both groups at T15.

Finally, the concentration of total salivary protein increased by 126% in EA and 438% in PAE, already showing a return to baseline levels at T5 for EA, but remaining with an increase of 213% in PAE, reaching negative levels in the T15 for EA and levels still high in PAE.

## 4. Discussion

Saliva plays a fundamental role in immunometabolism, food digestion, immune response, drug absorption, and taste. With age, salivary flow can decrease, resulting in malnutrition, recurrent respiratory and digestive tract infections, and a diminution in taste, with implications for the quality of life.^9^

Given that the continuous use of different pharmacological agents in this age group is common, for example in the form of polypharmacy (where three or more medications are taken), hyposalivation (a side-effect of many drugs) can be an issue.^10^

Age, nutritional status, hygiene, medication use, and previous illnesses can all affect salivary flow, but the present study has shown that the level of physical training and regular exercise can be added to the list. The decrease in salivary flow was almost double in PAE relative to AE; recovery was rapid in the second group but remained below baseline levels in the first. These findings corroborate those of a previous study on rats, where regular exercise suppressed all age-induced changes in salivary flow.^11^ We have also indicated that not only regular practice but the level of training also has an influence.

To confirm and give robustness to the findings, correlating salivary flow with physical conditioning, we indicate that cardiovascular parameters were inversely correlated with salivary flow; both groups showed a strong correlation with the double product but with a coefficient of almost double in PAE. It was also possible to observe that the change in this parameter was caused by systolic blood pressure in the AS group. In PAE, heart rate was the most closely correlated, with almost twice the correlation coefficient.

We also show that not only salivary flow presents itself differently according to the level of physical conditioning. The concentration of salivary nitric oxide, an essential mediator of intra- and extracellular processes mediating immunometabolism, and which even influences body composition,^12^ differed between the groups. An elevation of this biomarker in EA of 22% was observed at the time of exhaustion, with a reduction at later times. In PAE, the elevation in the TE was 34%. It continued to increase until it reached 94% of elevation at T5, followed by a reduction to the point where it was 27% at T15 relative to baseline values.

Salivary immunoglobulin A, an essential agent in the immunity of the mouth and respiratory and digestive tracts, and which is responsible for oral microbiome homeostasis,^13^ showed an increase of 37% in the TE for EA and only 7% in PAE. At T5, it continued upward in EA, reaching 41% at baseline levels, while PAE saw a 15% increase. Both saw a reduction in the final study time.

The concentration and activity of salivary amylase, an enzyme involved in the initiation of polysaccharide digestion and the modulation of immunity, can be altered by mental health,^14^ body composition, diet,^15^ and age.^16^ The present study has shown that it can also be affected by physical conditioning, even in elderly individuals: EA showed an increase of 130% at the time of exhaustion, with a subsequent reduction thereafter. By contrast, PAE showed a 68% increase in the time to exhaustion with a later peak (T5), reaching an elevation of 100%, and a subsequent sudden reduction. The results suggest that EA were more immunometabolically efficient.

Total salivary proteins, which have marked antifungal, antiviral, and antibacterial, pH homeostasis, digestive, mineralization, and tissue protection functions, are strongly influenced by age, dietary habits, and environmental factors.^17^ The present study has shown that they are also influenced by the level of training and physical conditioning. While EA showed a 126% increase in TE with a reduction to baseline values five minutes later, PAE showed an increase of 438% at the time of exhaustion, remaining high at T5 (213%) and T15 (47%).

Finally, antagonistic behaviors were observed between the groups regarding SNO with SBP and DBP, SA with DBP, and synergistically, but with notably different intensities for IgA-s with HR and DP, SA with SBP, and STP with HR. The impact of training levels for the age group under study has been revealed herein for the first time.

The unsupervised machine learning model was presented as a crucial exploratory tool to search for correlations between variables and the formation of homogeneous groups among themselves and heterogeneous groups. The present study used the Euclidean measure of dissimilarity to assess the distance between variables and multivariate cluster analysis with Classical Clustering (Agglomerative Hierarchical Method) and Nearest neighbor (single linkage), and sorting by Principal Component Analysis (PCA) and Correspondence Analysis (CA).

## 5. Conclusions

The level of physical conditioning in the 60+ age group has been shown to impact cardiovascular parameters, salivary biomarkers, and their correlation. Even when the two sub-groups in question regularly practiced the same type of exercise and competed with the same regularity, their level of training affected the behavior of the study variables.

Therefore, geriatricians, gerontologists, physical trainers, nutritionists, and other professionals who work with this age group must pay attention not only to relevant reference values but also training levels. They should also have respect for the longevity and quality of life of individuals at such a noble age.

EA – Elderly Athletes; PAE - Physically Active Elderly; BL - Blood Lactate; STP – Salivary Total Protein; SA - Salivary Amylase; IgA-s – Salivary Immunoglobulin A; SF - Salivary Flow; SNO - Salivary Nitric Oxide; DP - Double Product; SBP – Systolic Blood Pressure; DBP - Diastolic Blood Pressure; HR – Heart Rate.

EA: Elderly Athletes
PAE: Physically Active Elderly
BL: Blood Lactate
STP: Salivary Total Protein
SA: Salivary Amylase
IgA-s: Salivary Immunoglobulin A
SF: Salivary Flow
SNO: Salivary Nitric Oxide
DP: Double Product
SBP: Systolic Blood Pressure
DBP: Diastolic Blood Pressure
HR: Heart Rate

## Data Availability

All data generated or analyzed during the study are included in the published article. Raw data can be requested from the corresponding author.

## Ethical Approval

The present study was approved by the Human Research Ethics Committee of the Federal University of Mato Grosso, opinion number 5.716.414.

## Consent

After an initial lecture on the procedures, objectives, risks, and benefits given by those involved in the study, the participants completed the Free and Informed Consent Form (ICF). They were told that they could withdraw at any time and that they would be withdrawn if they did not follow the protocols.

## Conflicts of Interests

The authors declare no conflicts of interest.

## Practical applications

Gerontologists, geriatricians, nutritionists, physical education professionals, public managers, and other professionals who work with older people must take into account the age-related traditional reference values for biomarkers and cardiovascular parameters and recognize that the level of physical training in this group affects the data. Studies such as the present one can be used as a parameter for different decision-making.

## Author’s Contributions

ABJ, FSE and AMMN conceived and designed the study. PJVC, ABJ, LCOG, RJSB, MJSB, FSE and AMMN collected the data. PJVC, ABJ, LCOG, RJSB, MJSB, FSE and AMMN analyzed the data and wrote the manuscript. All authors read and provided critical feedback on the manuscript before approval.

## Acknowledgements

We thank the participants who agreed to participate in the study and the health professionals who directly or indirectly helped to make it all happen.

## Notes

### Competing Interest Statement

The authors have declared no competing interest.

